# Structural origins of altered spectroscopic properties upon ligand binding in proteins containing a fluorescent non-canonical amino acid

**DOI:** 10.1101/2021.04.24.441261

**Authors:** Patrick R. Gleason, Bethany Kolbaba-Kartchner, J. Nathan Henderson, Chad R. Simmons, Jeremy H. Mills

**Affiliations:** The Biodesign Center for Molecular Design and Biomimetics, Arizona State University, Tempe, AZ, 85287, USA; School of Molecular Sciences, Arizona State University, Tempe, AZ, 85287, USA

**Keywords:** Fluorescent Proteins, Non-canonical Amino Acids, X-ray Crystallography, Biosensors, Spectroscopy

## Abstract

Fluorescent non-canonical amino acids (fNCAAs) could serve as starting points for the rational design of protein-based fluorescent sensors of biological activity. However, efforts toward this goal are likely hampered by a lack of atomic-level characterization of fNCAAs within proteins. Here, we describe the spectroscopic and structural characterization of five streptavidin mutants that contain the fNCAA L-(7-hydroxycoumarin-4-yl)ethylglycine (7-HCAA) at sites proximal to the binding site of its substrate, biotin. Many of the mutants exhibited altered fluorescence spectra in response to biotin binding, which included both increases and decreases in fluorescence intensity as well as red or blue shifted emission maxima. Structural data were also obtained for three of the five mutants. The crystal structures shed light on interactions between 7-HCAA and functional groups—contributed either by the protein or substrate—that may be responsible for the observed changes in the 7-HCAA spectra. These data could be used in future studies aimed at the rational design of fluorescent, protein-based sensors of small molecule binding or dissociation.

## INTRODUCTION

Genetically encoded non-canonical amino acids (NCAAs) possess chemical functionalities that are not found among the twenty proteogenic amino acids.^1^ The side chains of some NCAAs are fluorescent and have emission wavelengths that fall in the visible range.^2–6^ Because such fluorescent non-canonical amino acids (fNCAAs) are translationally incorporated, they are not limited to placement at surface exposed residues or the protein termini, which are common limitations to other frequently used methods of fluorescently labeling proteins. Furthermore, most fNCAAs are similar in size to naturally occurring amino acids, which suggests that they can be incorporated in less accessible areas of a protein without a great risk of disruption of folding or function. Finally, many fNCAAs exhibit fluorescence spectra that are responsive to local environments.^2–4^ These features suggest that fNCAAs could represent excellent starting points for the development of novel, protein-based fluorescent biosensors.

Among the fNCAAs, L-(7-hydroxycoumarin-4-yl)ethylglycine^3^ (7-HCAA), has found extensive use in protein-based, fluorescent sensors that report on protein-ligand interactions,^7–9^ protein-protein interactions,^10–12^ enzyme-substrate binding,^13,14^ and changes in tyrosine phosphorylation state.^15^ A likely reason for 7-HCAA’s widespread use is that it possesses tunable functional groups that can be used to modulate its fluorescence output. For example, the 7-hydroxycoumarin (7-HC) moiety can exist in a number of tautomeric or ionic forms in the ground state that each give rise to distinct absorption/emission maxima (Scheme S1).^16^ Furthermore, 7-HC is a photoacid wherein the pK_a_ of the phenol drops from ∼7.8 in the ground state to ∼0.4 in the excited state (Scheme S1).^16^ When a proton acceptor (e.g., water) is present, the phenolic proton is rapidly shuttled from the excited hydroxycoumarin to the proton acceptor in a process termed excited state proton transfer (ESPT).^16^ If ESPT is blocked, the predominant emitting species is the phenol rather than the phenolate and the emission maximum is shifted from 450 nm to 380 nm (Scheme S1).^17–19^

The diverse chemical environments found in proteins could potentially be used to alter 7-HCAA emission, but only if appropriate sites of incorporation can be identified. In previous studies,^7–11,13–15^ two general approaches were used to select sites of 7-HCAA incorporation: The first approach relies on Förster Resonance Energy Transfer (FRET) ^7–9,11,12^ between 7-HCAA and a second fluorophore on the target protein. Although powerful, FRET-based techniques are best suited for use in proteins in which altered distances between the two fluorophores are expected as a consequence of conformational changes related to protein function. In a second approach, structure-based analyses were used to identify sites of 7-HCAA incorporation that were close to ligand binding sites,^13,14^ protein-protein interfaces,^10^ or a residue that was known to be post-translationally modified.^15^ Both approaches afforded protein-based sensors in which changes in 7-HCAA fluorescence could be used to monitor protein function; however, none of these 7-HCAA-modified proteins were structurally characterized in the initial studies. Recently, the structure of a 7-HCAA-containing sensor of protein-protein interactions was reported.^20^ However, this study was limited to a single protein and only one site of 7-HCAA incorporation. Without additional structural detail regarding the response of 7-HCAA to distinct chemical environments, it is difficult to envision how 7-HCAA might be systematically applied to address distinct challenges in the future.

In this study, we hoped to gain additional insight into how chemical environments affect the fluorescence properties of 7-HCAA. To achieve this, we genetically encoded 7-HCAA at five positions in the biotin-binding protein streptavidin (SAV) that were in close proximity to the substrate binding pocket. The photochemical properties of each mutant were assessed in the presence and absence of biotin, and biotin-dependent changes in fluorescence were observed in four of the five mutant proteins. Structural characterization of three of these mutants provided insight into how the protein environment surrounding 7-HCAA may alter its photochemical properties. The results of this study should prove useful in future efforts to develop novel protein-based fluorescent sensors of analyte binding.

## RESULTS AND DISCUSSION

### Selection of sites of 7-HCAA incorporation

Our primary goal in selecting potential sites of 7-HCAA incorporation was to identify residues that were likely to experience distinct chemical environments in the unbound (apo) and biotin-bound (holo) forms of SAV. We began by analyzing a high-resolution (0.95 Å) structure of SAV bound to biotin (PDB ID: 3ry2)^21^ and identified residues in close proximity to the biotin binding site. Because 7-HCAA contains a flexible ethylene linker between the peptide backbone and the 7-HC side chain (Figure S1), it is difficult to confidently predict the orientation that the 7-HCAA side chain would adopt within the folded SAV. However, we reasoned that substitution of 7-HCAA at sites suitably close to the substrate binding pocket could result in biotin-dependent alterations of the chemical environments in the vicinity of the fNCAA. Importantly, our approach was agnostic with respect to the nature of the changes in fluorescence that might occur in going from apo to holo SAV; rather, we simply hoped to identify sites where a biotin-dependent change in chemical environment seemed possible. Ultimately, we identified five residues surrounding the biotin binding pocket (L25, L110, S112, W120, and L124; Figure 1) that were not directly involved in hydrogen bonding interactions with biotin. Although the side chains of residues 45-50 are in close proximity to the biotin substrate, they fall on the flexible “loop 3/4”, which adopts “open” and “closed” conformations in the absence and presence of biotin, respectively; thus, these residues were excluded from consideration.^22^

**Figure 1.**
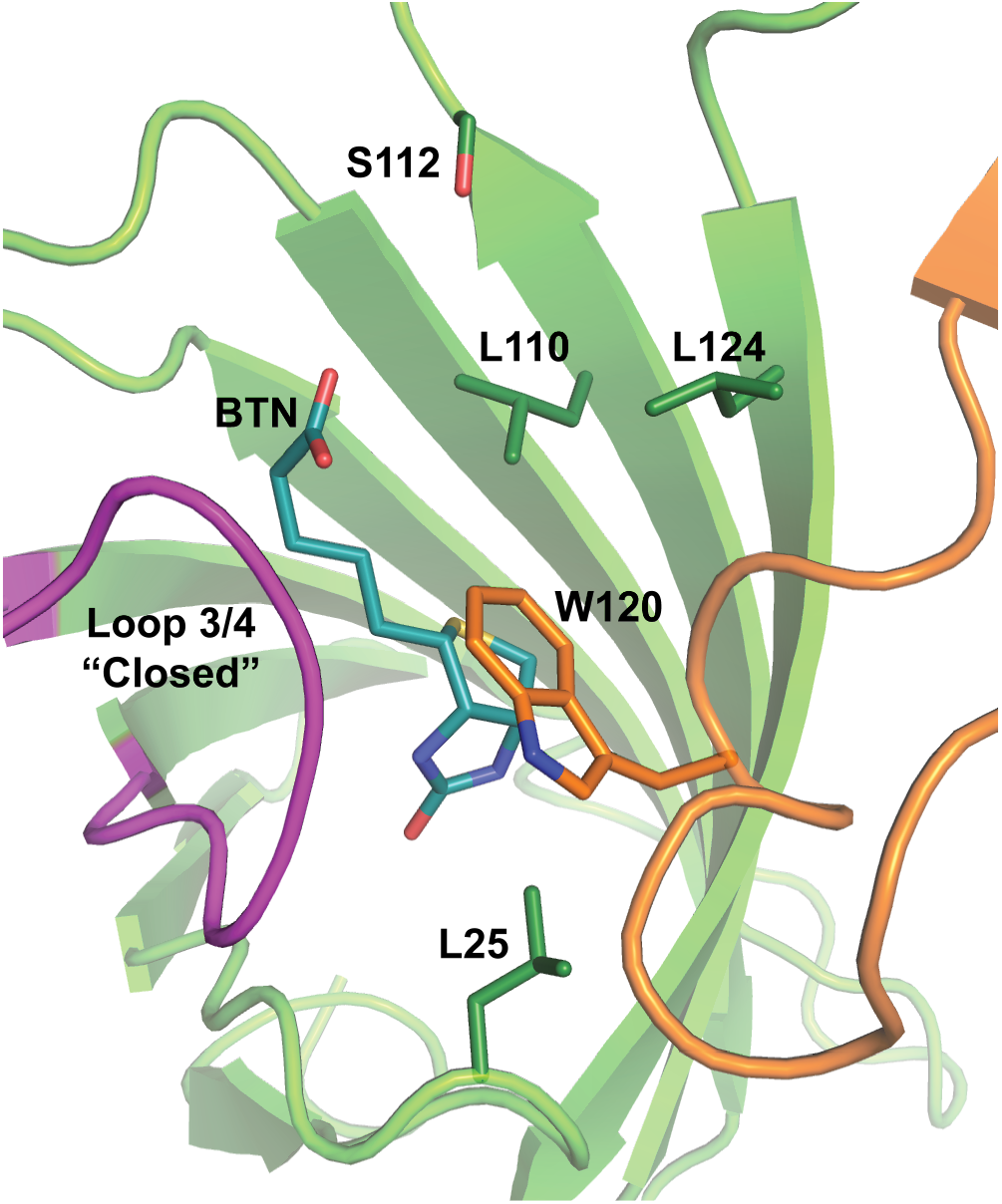
Sites of 7-HCAA incorporation in streptavidin. The structure of one subunit of the streptavidin tetramer (Green) and a portion of an adjacent subunit (Orange) are shown in cartoon form (PDB ID: 3ry2). Biotin (BTN; teal) and the side chains of the five residues that were substituted with 7-HCAA (dark green) are shown as sticks. Residue W120 (dark orange sticks) lies on an adjacent streptavidin subunit, but interacts with the biotin molecule in the subunit in green. Loop ¾ is shown in the closed position (dark purple).

### Spectroscopic and structural characterization of 7-HCAA containing SAV mutants

7-HCAA incorporation was achieved using the Amber codon suppression methodology developed by Schultz and co-workers.^3,23^ Full-length protein expression was observed for all five mutants only when 7-HCAA was present in the growth media (Figure S2a). Furthermore, fluorescent bands of the correct size were observed for all mutants, which confirmed 7-HCAA incorporation (Figure S2b). Mutant proteins were refolded from inclusion bodies using a modified version of a previously reported protocol^24^ and were further purified using immobilized metal ion, anion exchange, and size exclusion chromatography (see Experimental Procedures in the Supplemental Information for full expression and purification and refolding details).

In order to explore the effect of biotin binding on the spectral properties of 7-HCAA, we collected absorbance and fluorescence spectra of each SAV variant in the absence and presence of saturating amounts (100 µM) of biotin. At neutral pH, 7-HC exhibits two absorbance maxima at ∼325 nm and ∼360 nm that correspond to the neutral and anionic species, respectively. Because the p*K*_a_ of 7-HC is ∼7.8,^16^ both the phenol and phenolate species should have been present under the assay conditions (20 mM Tris-HCl, pH 7.0, 150 mM NaCl) unless the protein environment had altered the apparent p*K*_a_ of the 7-HC phenol. All proteins were excited at 340 nm, which corresponds to the isosbestic point of 7-HCAA. When possible, structural data were also collected for each protein variant. The spectroscopic and structural data from all mutants assayed in this study are described below.

#### L25X Characterization

The L25X mutant (where X signifies mutation of the native residue to 7-HCAA) shows a 40% increase in emission at 450 nm in the presence of biotin (Table 1, Figure 2b). A single absorbance maximum at 325 nm was observed for the L25X mutant in both the apo and holo forms (Figure 2b inset), which suggests that 7-HCAA exists primarily in the neutral form irrespective of the presence of biotin. Structural characterization of this mutant was precluded by low protein expression yields. However, the fact that the absorbance spectra of the apo and bound forms are nearly identical suggests that the chemical environment surrounding the 7-HC phenol in this mutant is not substantially altered by the addition of biotin.

**Table 1.** Spectroscopic data for SAV mutants compared with free 7-HCAA.

| Mutant | Abs. Max<br>(apo / holo) | $\Delta$ Abs. Intensity<br>(holo - apo) | Em. Max<br>(apo / holo) | $\Delta$ Em. Intensity<br>(holo - apo) |
| --- | --- | --- | --- | --- |
| 7-HCAA<sup>a</sup> | 326 | N/A<sup>b</sup> | 452 | N/ A<sup>b</sup> |
| L25X | 325 / 325 | -3% | 453 / 455 (+2 nm) | +40% |
| L110X | 324 / 324 | 12% | 449 / 435 (-14 nm) | -44% |
| S112X | 331 / 325 (-6 nm) | 10% | 451 / 455 (+4 nm) | +72% |
| W120X | 329 / 332 (+3 nm) | -18% | 453 / 433 (-20 nm) | +135% |
| L124X | N/A<sup>b</sup> | N/A<sup>b</sup> | 450 / 450 | +6% |

<sup>a</sup>Unincorporated L-(7-hydroxycoumarin-4-yl)ethylglycine in TBS (25 mM Tris-HCl, pH 7.0, 150 mM NaCl).
<sup>b</sup>Values were either not applicable or not obtainable.

**Figure 2.**
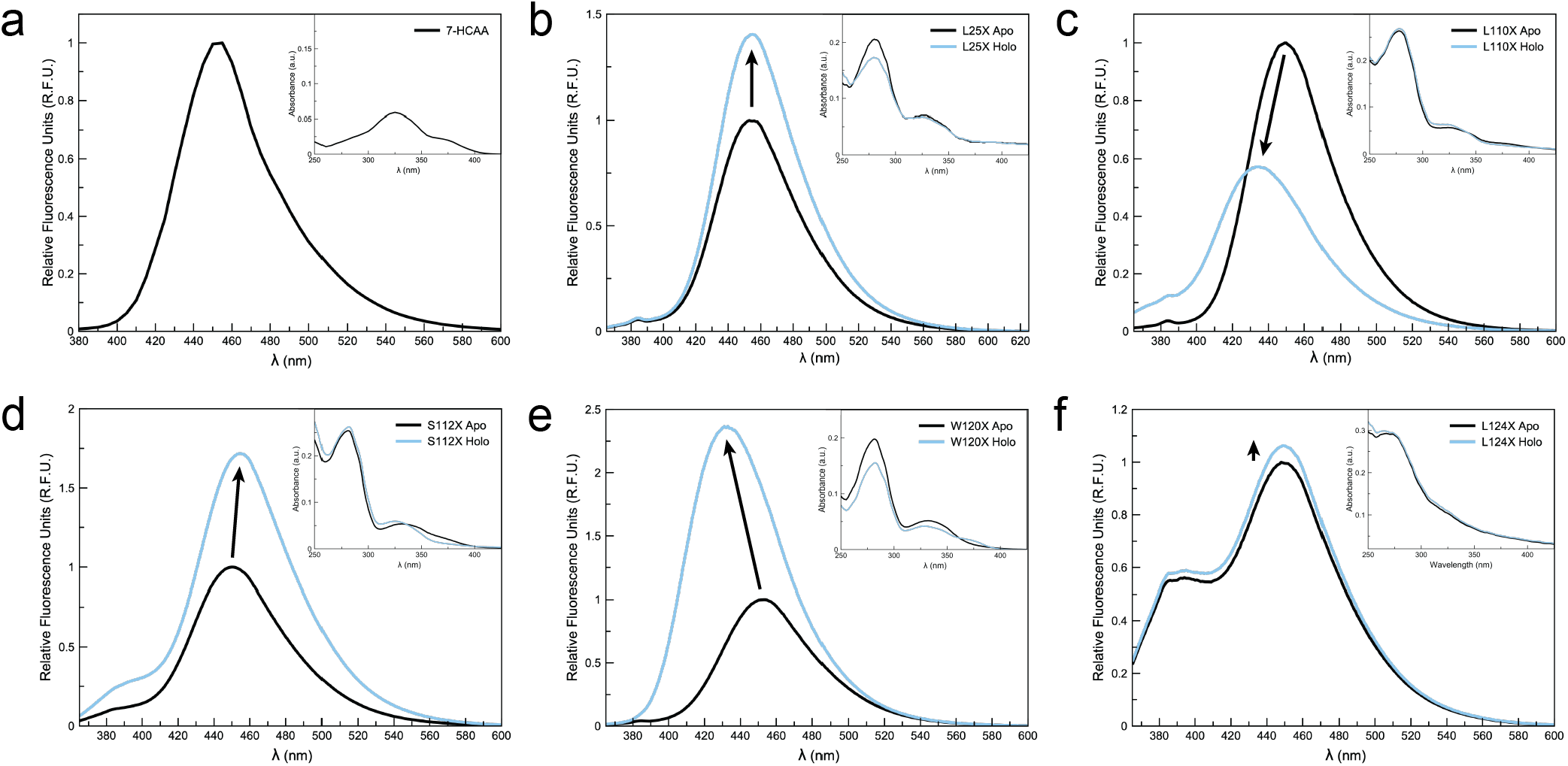
Normalized fluorescence and absorbance spectra of 7-HCAA (a), and the L25X (b), L110X (c), S112X (d), L124X (e), and W120X (f) mutants. Fluorescence spectra of the apo and holo forms of each mutant are shown in black and cyan lines, respectively. The fluorescence intensity of the apo form of each protein was set to unity in each case. Absorption spectra for each mutant are shown in the insets; black and cyan lines again represent the apo and holo forms of SAV, respectively. Studies were carried out at ∼10 uM protein with 100 µM biotin at an excitation wavelength of 340 nm; all spectra are the average of three independent measurements.

In an effort to gain additional insight into the orientation adopted by 7-HCAA in this mutant, we used the PyMOL molecular viewing software^25^ to mutate residue L25 to 7-HCAA in the apo form of SAV (PDB ID: 1swc). In this analysis, the ethylene linker of 7-HCAA was fit to the existing rotamer of the native leucine residue up to the γ-carbon (Figure S3) and rotation around χ_2_ and χ_3_ was sampled. Although we cannot be certain that 7-HCAA adopts this rotamer in the L25X mutant, this placement immediately suggested a number of residues with which 7-HCAA could interact in ways that are consistent with the spectroscopic data. First, a hydrogen bond could be formed between the 7-HCAA phenol and residue S45 (Figure S3a). Another possibility is the formation of hydrogen bonds between the 7-HCAA phenol and residues Y43 and D128 (Figure S3b). Hydrogen bonding interactions between 7-HCAA and surrounding residues have been suggested as a potential quenching mechanism in a previous study.^20^ If any of these interactions were to form, it is likely that they would be disrupted upon biotin binding, which could give rise to the observed increase in fluorescence in the holo form of this protein. Although purely conjectural, the analysis above highlights residues that may serve to quench 7-HCAA fluorescence when in proximity to the fNCAA.

#### L110X Characterization

The L110X mutant showed a 44% decrease in emission intensity and a 14 nm blue shift in emission maximum (from 449 nm to 435 nm) with the addition of biotin (Table 1, Figure 2c). The absorption spectra of both the apo and holo forms of L110X show maxima at ∼320 nm (Figure 2c inset). The apo form also shows a slight shoulder at ∼360 nm, which is indicative of a mixed population of neutral and anionic 7-HCAA. However, the population of the neutral species (λ_max_ = ∼320) increases by ∼33% upon substrate binding. This observation suggests the possibility that new interactions between the protein and/or biotin are forged in the holo form that might induce partial protonation of 7-HCAA.

To explore the origin of the fluorescence change, we solved the crystal structures of the L110X mutant in both the absence and presence of biotin to 1.55 Å and 2.10 Å resolution, respectively. In the apo form (Figure 3a), 7-HCAA extends into the biotin-binding pocket, and the 7-HC side chain is in van der Waals contact with residues W120 and L124; the C_β_ atoms of S88 and S112 pack against the opposite face of the coumarin ring. In the holo structure, clear electron density is observed for the biotin molecule (Figure 3b) and alignment of this structure with a previously reported, high resolution structure of biotin-bound SAV (PDB ID: 3ry2) indicated that very little structural rearrangement had occurred due to the 7-HCAA substitution (Figure S4). Namely, when all binding site residues within 6 Å of 7-HCAA are considered, the calculated all-atom root-mean-squared deviations (RMSDs) are <0.5 Å in all subunits. The maintenance of the positions of the residues within the binding site allows biotin to bind in its native orientation; superimposition of the structures of the holo forms of the L110X mutant and wild type SAV indicate an RMSD of 0.266 between the two biotin molecules. Furthermore, loop 3/4 is in the closed configuration in all subunits of L110X in the solved structure, which suggests that the placement of 7-HCAA within the binding site does not preclude the large structural rearrangements that occur upon biotin binding.^21,22^

**Figure 3.**
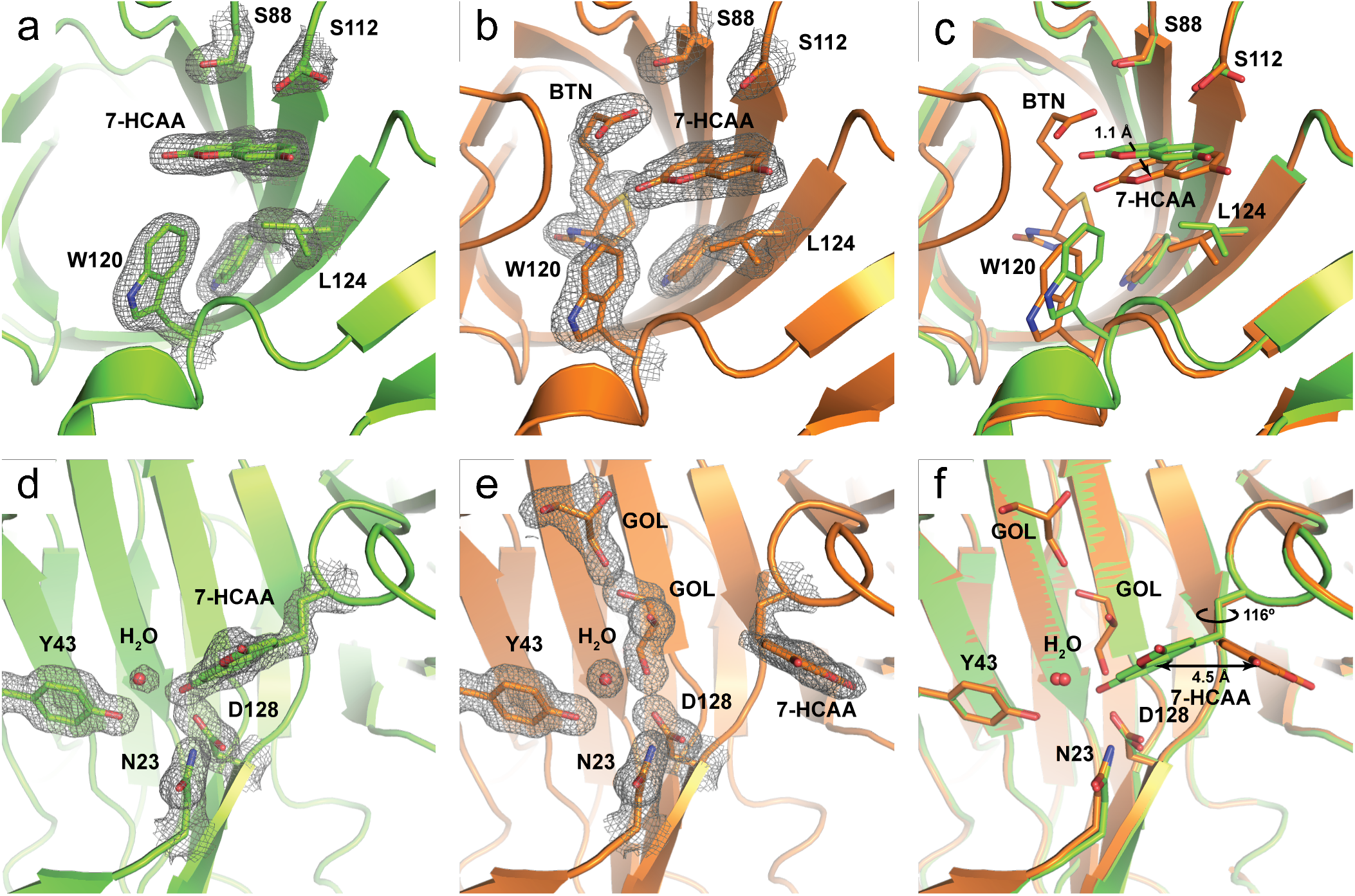
Crystal structures of the 7-HCAA substituted streptavidins. Structural data for the L110X mutant in the apo and holo forms are shown in panels a and b, respectively. An overlay of the apo (green) and holo (orange) forms of the L110X mutant are shown in panel c. Structural data for the W120X mutant in the apo form is shown in panel d. Panel e shows the W120X mutant in the presence of glycerol, which had been used as a cryo-protectant. An overlay of the apo (green) and holo (orange) structures of the W120X mutant is shown in panel f. 7-HCAA, biotin (BTN) and glycerol (GOL) and residues in the vicinity of 7-HCAA are shown as sticks. Electron density around each residue is shown as a 2FO-FC map contoured to 1 σ.

In the bound structure (Figure 3b), biotin directly interacts with the 7-HCAA side chain, primarily through van der Waals interactions between the carboxylic acid on biotin and the pyrone ring of the 7-HC. This interaction also induces a slight (∼1.1 Å) re-orientation of the 7-HC side chain (Figure 3c), which is accommodated by a 17.1° alteration of the C_α_-C_β_-C_γ_ angle of L124 (114.9° in the apo form vs 132.0° in the holo form) that is not seen in other holo structures of SAV (e.g., PDB ID: 3ry2). This causes adjacent subunits of the SAV tetramer to also shift 1.1 Å in relation to each other in order to avoid clashes between L124 on the same subunit and K121 on an adjacent subunit.

Our efforts to rationalize the biotin-dependent change in fluorescence for this mutant focused mainly on the distinct chemical environments surrounding 7-HCAA in the apo and bound states. Chief among these appears to be the close packing between the negatively charged biotin carboxylate and 7-HCAA. In previous work on the “fruit” series of fluorescent proteins, red- and blue-shifted emission maxima were observed as a consequence of altered electrostatic environments in the vicinity of the chromophore.^26^ In a related study, red shifted emission maxima were observed for the cyan fluorescent protein amFP486 when either of two negatively charged glutamates in close proximity to the chromophore were mutated to neutral glutamine residues.^27^ The interaction between the biotin carboxylate and 7-HCAA, which, like the chromophore in amFP486, is a photo acid, represents the opposite of the aforementioned result. Namely, a 14 nm blue shift in fluorescence maximum is observed in this study when a negatively charged functional group is introduced in the vicinity of 7-HCAA.

Although these literature precedents provide a potential explanation for the observed blue shift in 7-HCAA fluorescence, the origin of the substantial decrease in emission intensity in the biotin-bound protein is less readily explained. Because 7-HCAA moves only slightly as a consequence of biotin binding, the chemical environments experienced in the apo and holo forms are highly similar save for the novel interaction with a carboxylic acid on the biotin substrate. This suggests that the presence of the biotin is also likely responsible for the observed quenching of fluorescence. The mechanism through which this occurs (e.g., collisional quenching or photoinduced electron transfer) is less clear and elucidation of the specific quenching mechanisms underpinning this phenomenon is beyond the scope of this work.

#### S112X characterization

The S112X mutant exhibited a 71% increase in fluorescence intensity upon biotin binding (Figure 2d). Furthermore, a notable shoulder at ∼380 nm is observed in the emission spectrum of the holo form that could arise from interactions that block ESPT in a population of 7-HCAA. Despite repeated attempts, we were unable to obtain crystals of the apo S112X mutant. However, a structure of the S112X mutant in the holo form was solved to 1.8 Å resolution; both the biotin and 7-HCAA are fully resolved in this structure (Figure S5).

Consistent with the other structures reported in this study, the position of the biotin in this structure is identical to that observed in the wild type protein (Figure S6), which suggests that the addition of 7-HCAA at position S112 did not substantially disrupt the native biotin binding interactions. While density for the 7-HCAA is clearly visible in both monomers of the asymmetric unit, it is weaker than the density of the surrounding residues. Furthermore, the 7-HCAA adopts two distinct conformations that are rotated 108° around the C_α_-C_β_ vector. In one of the monomers (chain A), the 7-HCAA is involved in a crystal contact via a hydrogen bond between the 7-HC phenol and the backbone carbonyl of Y83 (Figure S7a). In the second subunit (chain B), 7-HCAA adopts a conformation that places its phenol in a reasonably hydrophobic environment generated by the side chains of L124 in the same subunit and L124 on an adjacent subunit (Figure S7b).

Because 7-HCAA in chain A is involved in a crystal contact, it is unlikely that this orientation in adopted in solution. We therefore cannot draw conclusions regarding the effects of this chemical environment on 7-HCAA fluorescence. In chain B, no solvent molecules are observed within the relatively hydrophobic environment surrounding the 7-HC side chain, which suggests that a neutral form of the phenol would likely be favored. This is consistent with the increase in 325 nm absorbance that occurs as a consequence of biotin binding as well as the slight increase in 380 nm emission that is observed upon biotin-binding. The predominant 450 nm emission that is observed for both the apo and holo forms of the S112X mutant is less readily explained by our crystallographic data. It is therefore likely that the orientation of 7-HCAA in chain A that is adopted in solution is responsible for the observed fluorescence spectra.

#### W120X characterization

The emission intensity of W120X increases 135% upon substrate binding and represents the largest biotin-dependent change in fluorescence observed in this study. Similar to the L110X mutant, a blue shift in emission maximum from 453 nm (apo form) to 433 nm (holo form) is evident (Table 1, Figure 2e).

We initially solved a structure of the apo W120X mutant to 1.17 Å resolution at pH 3.5. No significant electron density was observed for the 7-HCAA; however, the presence of positive electron density near residue D128 suggested the possibility that a population of 7-HCAA was interacting with D128 through a hydrogen bond. We rationalized that the low pH of the crystallization conditions may have produced a significant population of protonated D128 that would preclude the formation of this hydrogen bond. We therefore soaked crystals obtained at pH 3.5 in buffers with increasing pHs up to a maximum of 6.0. We ultimately collected diffraction data to 1.55 Å resolution from a crystal that had been soaked to pH 5.5; electron density for 7-HCAA is clearly visible in this structure (Figure 3d). Notably, an anomalously short (< 2.4 Å) hydrogen bond between the phenolic oxygen of 7-HCAA and the carboxylate of residue D128 was observed in both chains of the asymmetric unit (Figure S8). Additional hydrogen bonding interactions were identified between the phenol of 7-HCAA, an ordered water molecule, and the phenol of tyrosine 43 (Figure S8).

Several attempts were made to either co-crystallize W120X with biotin or to soak the substrate into the apo W120X crystals, but these efforts were unsuccessful. However, in a previously solved structure of the W120X mutant (1.50 Å resolution, pH 5.5) where glycerol was used as a cryoprotectant, two glycerol molecules were observed in the biotin binding pocket (Figure 3e). When superimposed with the 3ry2 structure, the bound glycerol molecules directly overlap with a number of atoms in the biotin molecule (Figure S9), which served to displace the 7-HCAA from the biotin binding pocket. This displacement is primarily a consequence of a substantial (116°) rotation about χ_2_ of 7-HCAA (Figure 3f). A spectrum of the W120X mutant in the presence of 50% (v/v) glycerol showed an 11% increase in fluorescence at 450 nm, which supports the notion that displacement of 7-HCAA from the binding pocket is primarily responsible for the observed change in fluorescence. Interestingly, the blue shift in emission maximum that was observed after the addition of biotin was not as pronounced in the presence of glycerol (Figure S10). One possible explanation for this is that a functional group on the biotin molecule is responsible for the shift in emission maximum that is observed in holo W120X. Additionally, we note that loop 3/4, which closes over the binding site when biotin binds, is in the open position in the glycerol bound structure. Any interactions between this loop and the 7-HCAA that are present in the holo form of SAV would not be apparent in this structure.

With respect to the observed fluorescence quenching in this mutant, it seems likely that the particularly short hydrogen bond observed between the 7-HCAA phenol and D128 in the adjacent SAV monomer (Figure S8) could be responsible. Similarly short hydrogen bonds have been observed in proteins between acidic residues and conjugated phenols, including 7-HCAA.^20,28,29^ It has previously been suggested^20^ that a short hydrogen bond between an acidic residue and the 7-HCAA could be responsible for quenching via backward transfer of a proton.^30–34^ Because this hydrogen bonding interaction is disrupted in the glycerol-bound structure of W120X, we believe that this interaction is likely responsible for the quenching observed in the apo state.

#### L124X characterization

Both the absorbance and emission spectra of the L124X mutant exhibit features distinct from those observed in all other variants. The absorbance spectra of both apo and holo L124X are essentially linear from 300 nm to 450 nm. The emission spectrum of this mutant exhibits a major peak at 450 nm with a significant shoulder at 380 nm. The substantial emission at 380 nm suggests that that ESPT from 7-HCAA could be at least partially blocked in this mutant. Although increases in emission intensity are observed at both 380 nm and 450 nm upon biotin binding, the magnitude of this change is substantially smaller than those observed for all other variants (Table 1, Figure 2). Unfortunately, L124X suffered from low initial expression yields and exhibited consistent decreases in yield throughout each stage of the purification process; characterization of this protein was limited to spectroscopic analysis alone. In an effort to better understand these data, we modeled 7-HCAA in place of residue 124 in the wild type apo structure of SAV (PDB ID: 1swc) using the PyMOL software. We immediately recognized that position 124 is directly adjacent to the identical residue in a symmetry related subunit; the C_α_—C_α_ distance between these residues is only 5.0 Å. Without structural data, it is difficult to confidently predict the orientations adopted by the two 7-HC moieties in this protein. However, given the close proximity of the sites of incorporation, the possibility exists that the two fluorophores would directly interact, which could potentially lead to the unique spectra we observed.

### Affinity of the streptavidin mutants for biotin

Structural characterization of three of the five SAV mutants indicated that the introduction of 7-HCAA had not caused substantial rearrangement of residues within the biotin binding pocket. However, it seemed likely that the presence of 7-HCAA within the substrate binding site could have altered the affinity of our mutant proteins for biotin. To test this, we measured the apparent dissociation constants for each of the stable SAV mutants (L110X, S112X, and W120X) using changes in the fluorescence intensity of 7-HCAA that occurred upon biotin binding. For S112X, the increase in absorbance at 450 nm was plotted against biotin concentration and an apparent *K*_*d*_ was established by non-linear regression (Table S1, Figure 4). For the L110X and W120X mutants, biotin binding resulted not only in a decrease and an increase in fluorescence emission intensity, respectively, but also showed a blue shifted emission maximum. We therefore plotted the ratio of 450 nm to 438 nm emission (the emission maxima of the apo and holo forms, respectively) against biotin concentration and derived apparent *K*_*d*_s using non-linear regression (Table S1, Figure 4). Apparent *K*_*d*_s of 568.6 nM for L110X, 1.6 µM for S112X and 6.2 nM for W120X were ultimately measured. Because the reported *K*_*d*_ of SAV for biotin is in the picomolar to high femtomolar range,^35^ it appears that the introduction of 7-HCAA resulted in a substantial loss in affinity relative to the wild type protein in all mutants for which apparent *K*_d_s were determined; however, reasonable affinities for biotin were still maintained. Nevertheless, the fact that a range of affinities was observed among the mutants suggests a direct path to the development of fluorescent sensors of small molecule metabolites that could function over a broad range of concentrations.

**Figure 4.**
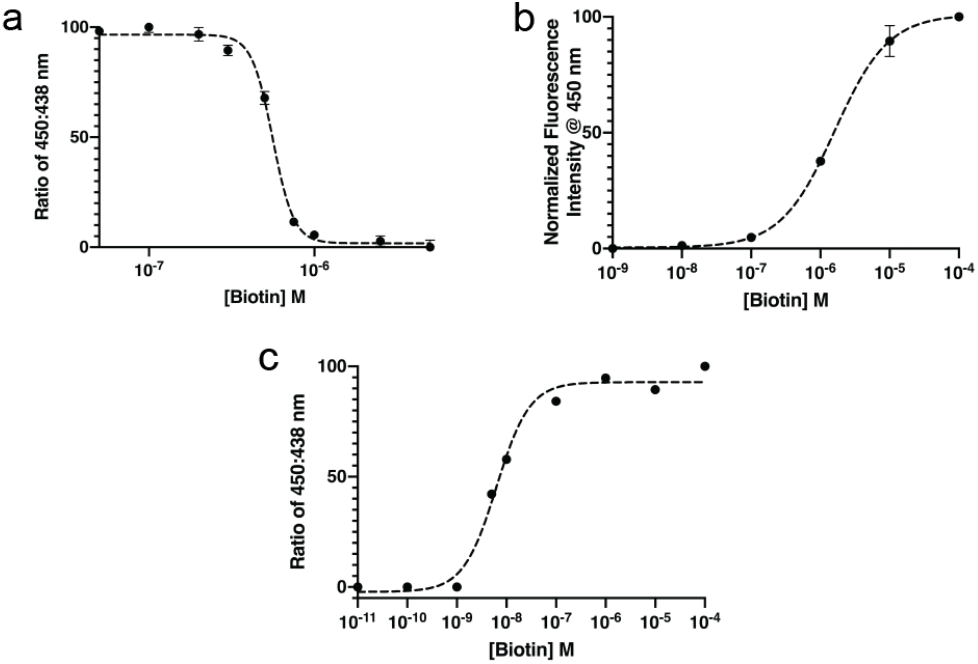
Biotin binding analysis for the L110X (a), S112X (b), and W120X (c) SAV mutants. For the L110X and W120X mutants, the ratio of 450 nm to 438 nm emission is plotted against biotin concentration. For the S112X mutant, the emission intensity at 450 nm is plotted directly against biotin concentration. Some error bars are too small to be visible.

### Implications of this study on the design of new fluorescent sensors

The primary goal of this study was to gain a deeper understanding of how protein environments can alter the fluorescence properties of 7-HCAA. Here, we summarize some of our findings and relate them to future rational design efforts aimed at generating new fluorescent sensors of metabolite binding.

One result of note is the ease with which fluorescent sensors of small molecule binding can apparently be developed using 7-HCAA. We employed a simplistic approach in which residues surrounding a small molecule binding site were mutated to 7-HCAA without further consideration of the local environments surrounding these sites. The fact that substrate-dependent changes to the fluorescence properties of 7-HCAA were observed in four of the five mutants suggests the versatility of this approach. Additionally, substituting 7-HCAA for native residues did not appear to substantially alter the positions of the side chains of amino acids in its vicinity and did not preclude substrate binding in any of the fully characterized proteins.

Collectively, these data suggest that significant flexibility exists with respect to potential sites of 7-HCAA incorporation, and that changes in its fluorescence properties are likely if they are in proximity to the binding pocket. A caveat to this observation is that substitution of 7-HCAA at sites that are close to symmetry-related residues in multimeric assemblies might result in unanticipated interactions of the 7-HC fluorophores. The fact that biotin-SAV interactions are among the strongest non-covalent interactions known was likely beneficial to this study; it is unclear if other proteins with lower affinities would be as amenable to modification as SAV. Nonetheless, the ease with which modulation of 7-HCAA’s fluorescent properties can be achieved using this approach is encouraging.

The structural data provide atomic-level portraits of 7-HCAA in a variety of chemically distinct environments and could therefore lead to the rational design of new fluorescent, protein-based sensors. We not only demonstrate that the fluorescence properties of 7-HC can be tuned by the surrounding environments, but also provide support for the mechanisms through which these spectroscopic changes may occur. For example, significant (>10 nm) blue-shifts in emission maximum were observed in two proteins (L110X and W120X) and structural characterization of one of these mutants (L110X) suggests that the electronic nature of the environment surrounding the 7-HC moiety is likely responsible. Furthermore, it is notable that interactions between 7-HCAA and the ligand itself can modulate 7-HC fluorescence. This suggests that 1) direct interactions between the fNCAA and the ligand can occur without ablating binding affinity and 2) that interactions between the functional groups contained within the ligand and 7-HC should be considered when identifying potential sites of 7-HCAA incorporation.

We also observed two interactions that appear to favor quenched states of 7-HCAA. In W120X, we attribute quenching to the presence of a short hydrogen bond that only appears to exist in the apo protein. Given the ease with which this hydrogen bond was identified in this study, it could be sufficient to bias sites of 7-HCAA incorporation to regions of target proteins that are both proximal to the substrate binding site and also contain acidic amino acids. Even if these criteria are not readily met, our results suggest the possibility of rationally engineering mutant proteins in which similar hydrogen bonds are likely to occur.

Another apparent mode for 7-HCAA quenching was observed in the L110X mutant, in which van der Waals contacts between the biotin substrate and 7-HCAA side chain appear to have quenched 7-HC fluorescence. While we believe that the presence of a negatively charged carboxylate group on biotin may have caused a blue shift in fluorescence in this mutant, it is possible that neutral portions of other substrates could quench, but not shift 7-HCAA fluorescence if bound in close proximity to the fluorophore.

Finally, in one instance, we observed a conformation of 7-HCAA that appears to have placed the phenol within a hydrophobic environment that at least partially favored the neutral form of 7-HC in both the ground and excited states (scheme S1). Because emission from the protonated form of 7-HC occurs in the UV range (380 nm), this principle could potentially be used to develop fluorescent sensors in which substrate binding is detectable by eye and therefore would not require the use of sophisticated instrumentation.

## CONCLUSION

Despite a wealth of fluorescent tools that have proven useful in the study of biological systems, the ability to develop fluorescent sensors of protein-small molecule interactions remains a challenge. Fluorescent non-canonical amino acids possess a number of properties (e.g., small sizes and versatility with respect to sites of incorporation) that may offer advantages for use in these studies. However, a lack of characterization of fNCAAs in proteins has likely limited their use. In this study, we incorporate the fNCAA L-(7-hydroxycoumarin-4-yl)ethylglycine into the biotin-binding protein streptavidin and demonstrate its ability to report on substrate binding events. Major advantages of this method relative to others (e.g., those in which FRET plays a central role) are that only a single fluorophore is required to generate a fluorescent sensor and that no conformational change in the protein is required to sense small molecule binding. Structural characterization of three SAV mutants provided an atomic-level understanding of the manner in which protein environments respond to 7-HCAA incorporation and also reported on the manner in which changes in local environments affect its fluorescence properties. In addition to demonstrating that 7-HCAA is a highly versatile, genetically encodable fluorophore, the spectroscopic and structural characterizations reported herein could help guide future efforts to rationally design new fluorescent sensors of protein-ligand interactions.

## Supporting information

Coordinate and Reflection Files

## Accession Codes

6udb, 6udc, 6ud1, 6ud6, 6uc3

## Notes

The authors declare no competing financial interest.

## Author Contributions

JHM conceived of the project. PRG and JHM designed the protein constructs and experiments. PRG and BKK performed molecular cloning, protein expression and protein purification. PRG, BKK, and JHM analyzed fluorescence data. PRG, under the guidance of JNH and CRS, crystalized proteins. PRG, JNH, and CRS collected and analyzed diffraction data. PRG and JHM drafted the manuscript. All authors have reviewed and commented on the final draft of the manuscript.

## ACKNOWLEDGMENT

The Berkeley Center for Structural Biology is supported in part by the Howard Hughes Medical Institute. The Advanced Light Source (ALS) is a Department of Energy Office of Science User Facility under Contract No. DE-AC02-05CH11231. Results derived from work performed at Argonne National Laboratory ANL, Structural Biology Center (SBC) at the Advanced Photon Source (APS) were supported under U.S. Department of Energy, Office of Biological and Environmental Research contract DE-AC02-06CH11357. Use of the Stanford Synchrotron Radiation Lightsource, SLAC National Accelerator Laboratory, is supported by the U.S. Department of Energy, Office of Science, Office of Basic Energy Sciences under Contract No. DE-AC02 76SF00515. The SSRL Structural Molecular Biology Program is supported by the DOE Office of Biological and Environmental Research, and by the National Institutes of Health, National Institute of General Medical Sciences (P30GM133894). The contents of this publication are solely the responsibility of the authors and do not necessarily represent the official views of NIGMS or NIH.

