## Supporting Information for "Structural origins of altered spectroscopic properties upon ligand binding in proteins containing a fluorescent non-canonical amino acid"

### Experimental Procedures

**Protein Expression & Purification:** Wild-type core streptavidin[1] (SAV) and DNA oligonucleotides for making each Amber stop codon mutation were ordered from Integrated DNA Technologies, Inc. The gene encoding streptavidin was cloned into a pET29b vector using Gibson assembly. Mutations were made via overlap extension PCR and Gibson assembly. *E. coli* BL21 Star (DE3) cells were transformed with both a pEVOL plasmid[2] containing a chloramphenicol resistance marker, two copies of the evolved CouRS synthetase, and an evolved tRNA specific to the evolved CouRS and a pET29 plasmid containing a kanamycin resistance marker and genes encoding the mutant streptavidins.

A single colony was used to inoculate 5 mL of 2xYT media, which was grown overnight in a shaking incubator at 37 ºC with 250 r.p.m. agitation until an $OD_{600}$ of ~5.0 was reached. The cells were centrifuged for 10 minutes at 4200x g and resuspended in fresh 2xYT media. Arabinose (0.2% w/v) and the non-canonical amino acid, 7-HCAA (1 mM), were added to each culture and incubated for 1 hour at 37 ºC with 250 r.p.m. agitation in order to express and activate the CouRS synthetase. Isopropyl-β-D-thiogalactoside (IPTG) was then added to a final concentration of 1 mM in order to induce expression of the SAV mutants. The cultures were then incubated between 8 and 16 hours at 30 ºC, with 180 r.p.m. shaking. Cells were harvested via centrifugation and lysed in cell lysis buffer (25 mM Tris-HCl, 0.1% Triton-X 100, 3 mM β-mercaptoethanol). Inclusion bodies were collected via centrifugation at 20,000 xg for 30 minutes and washed 3-4 times with wash buffer (25 mM Tris-HCl, 150 mM NaCl, 0.1% Triton-X 100). The washed inclusion bodies were then solubilized with 6M Guanidine-HCl. Solubilized protein was refolded in refolding buffer (25 mM Tris-HCl, 20 mM Imidazole, 500 mM NaCl) by adding the protein solution dropwise to the fastest part of a rapidly spinning solution of refolding buffer.[3]

The streptavidin mutants contained a C-terminal 6x His-tag that allowed purification on a nickel-NTA resin (HisTrap FF, GE Healthcare). After loading onto the column, contaminant proteins were removed by flowing five column volumes (CV) of Ni-NTA wash buffer (25 mM Tris-HCl pH 8.0, 20 mM Imidazole, 500 mM NaCl) over the column followed by elution of SAV with five CV Ni-NTA elution buffer (25 mM Tris-HCl pH 8.0, 500 mM Imidazole, 150 mM NaCl). Fluorescent fractions were collected, combined, diluted 10x in IEC wash buffer (25 mM Tris-HCl pH 8.0, 10 mM NaCl) and purified further on an anion exchange column (HiTrap Q FF, GE Healthcare). The column was washed with five CV of IEC wash buffer followed by a mixture of 90% wash buffer and 10% elution buffer (25 mM Tris-HCl pH 8.0, 500 mM NaCl). Proteins were then eluted with five CV of 100% IEC elution buffer. Again, fluorescent fractions from the IEC purification were collected and combined. The combined fractions were then concentrated using a 10 kDa molecular weight cutoff spin column (Amicon) to 500 µl. Concetrated protein was injected onto a size-exclusion column (Superdex 200 increase 10/300 gl, GE Healthcare) and eluted using Sizing Buffer (25 mM Tris-HCl pH 7.0, 150 mM NaCl). Fractions that were fluorescent were collected, consolidated, and concentrated to a volume of ~1 mL on a 10 kDa molecular cutoff spin column (Amicon). Fractions were then refrigerated at 4 ºC for use in experimental procedures.

**SDS-PAGE analysis of mutants:** Single colonies of *E. coli* harboring each SAV mutant were used to inoculate 5 mL cultures of 2xYT medium supplemented with 50 µg/mL kanamycin and 34 µg/mL chloramphenicol. Cells were grown overnight at 37 ºC with 250 r.p.m. shaking. The cultures were then centrifuged at 4,200 xg. and the media was decanted. Each pellet was resuspended in 5 mL of fresh 2xYT media containing 0.2% arabinose, 50 µg/mL kanamycin and 34 µg/mL chloramphenicol. The cultures were divided equally and 7-HCAA (1 mM) was added to only one of the two cultures. The cultures were then incubated at 37 ºC and 250 r.p.m. shaking for 1 hour. IPTG was then added to a final concentration of 1 mM and protein expression was allowed to continue at 30 ºC with 225 r.p.m. shaking for 16 hours. 15 µl of each culture or 1 µg of commercially available Streptavidin (Sigma) were mixed with 15 µl 4x Laemmli loading buffer and was incubated at 95 ºC for 10 minutes followed by 25 ºC for 5 minutes. 15 µl of each sample was loaded into each well of a 4% stacking / 15% resolving polyacrylamide gel and subjected to 125 V for 90 minutes.

**Western Blotting:** An acrylamide gel was equilibrated in 25 mL of transfer buffer (25 mM Tris, pH 8.3, 192 mM glycine, 20% methanol, 0.1% SDS) for 60 minutes. Protein from the acrylamide gel was transferred to a nitrocellulose membrane using a semi-dry transfer apparatus (Bio-Rad) until the pre-stained ladder (Precision Plus Protein, New England Biolabs) was completely transferred to the membrane (30 minutes at 25 V). The membrane was removed from the apparatus and incubated with a TBS-T blocking solution (25 mM Tris, pH 8.0, 150 mM NaCl, 0.1% v/v Tween-20, 5% w/v powdered nonfat milk) for 60 minutes. The membrane was then washed 3 times for 5 minutes with 25 mL of TBS-T (25 mM Tris, pH 8.0, 150 mM NaCl, 0.1% v/v Tween-20) on an orbital shaker, followed by incubation in TBS-T with an HRP-conjugated anti-streptavidin antibodies (Abcam; 1:10000 dilution) at room temperature for 1 hour on an orbital shaker. The membrane was then washed with 25 mL TBS-T 3 times for 5 minutes each on an orbital shaker. A 1x solution of DAB solution (Pierce) was created immediately before use by adding 2.5 mL of 10x DAB with 22.5 mL of Stable Peroxide Substrate Buffer. Finally, the membranes were incubated for 30 minutes in the 1x DAB solution, air-dried, and imaged.

**Spectroscopic Analysis:** All spectroscopic experiments were performed in a Tris-Buffered Saline (TBS) solution (25 mM Tris-HCl pH 7.0, 150 mM NaCl) using a 1 cm quartz cuvette (Starna Cells). Each apo mutant was concentrated to 100 µl and then diluted to a final absorbance of 0.05 at 340 nm. The samples were then split and either biotin (100 µM final concentration, dissolved in 30% DMSO) or TBS was added to each. A SpectraMax M5 spectrophotometer was used to take absorbance spectra from 200 nm to 750 nm at 1 nm intervals in triplicate. A Horiba Nanolog fluorimeter was used to measure relative fluorescence intensities. Both the excitation and emission slit widths were set to 2 nm and the spectra were taken from 365 nm to 625 nm in triplicate with excitation at 340 nm.

**Crystallization:** A CrystalMation Phoenix (Rigaku) was used to screen a 10 mg/mL solution (10 mM HEPES, pH 5.5, 75 mM NaCl) of both apo and holo L110X, S112X, and W120X against three sitting drop vapor diffusion crystal screening libraries (Hampton Research, Crystal HT, Index HT, and PEG/Ion HT). Each screen contains 96 conditions, and each condition was tested twice (v/v ratios of protein to reservoir drop) for a total of 576 total conditions per protein in 200 and 300 nl drop sizes. Conditions that produced crystals after 5 days were then recapitulated using larger volume sitting drop vapor diffusion. Streptavidin L110X apo and holo crystals were grown with a reservoir solution of 0.1 M Bis-Tris, pH 6.5, 25% w/v polyethylene glycol 3350. Streptavidin S112X holo, and W120X apo crystals were grown with a reservoir solution of 0.1M citric acid, pH 3.5, 3.0 M NaCl. Each drop contained 2 µL of protein solution mixed with 2 µL of reservoir solution, crystals were grown until no new growth was visible (approximately 1 week). Both the S112X and W120X crystals were soaked in a new reservoir solution of 0.1M citric acid, pH 5.5, 3.0 M NaCl. Three 2 µL drop exchanges were performed over the course of 12 hours and the crystals were allowed to soak overnight. The cryoprotectant was 25% PEG 3350 for L110X, S112X and the apo crystal for W120X. 25% glycerol was used as cryoprotectant for the W120X glycerol bound crystal.

**Data Collection and Structure Determination.** Diffraction data from the L110X apo and W120X apo crystals were collected at the Berkeley Center for Structural Biology (BCSB) from the Advanced Light Source (beamline 5.0.2) on a Dectris Pilatus3 6M detector. Data from the L110X holo and W120X glycerol bound crystals were collected at the Argonne National Laboratory Advanced Photon Source (beamline 19-ID) on a Dectris Pilatus 6M detector. Diffraction data from the S112X holo crystals were collected at the Stanford Synchrotron Radiation LightSource (beamline BL9-2) on a Dectris Pilatus 6M detector.

Crystals were flash frozen in liquid nitrogen prior to data collection at 100 K. Datasets for the L110X and W120X crystals were indexed, refined, integrated, and scaled using the HKL-3000 software package. The S112X dataset was indexed, refined and integrated using Mosfilm and scaled using Pointless. Structures were then solved by molecular replacement using Phaser19 using an all glycine model of SAV with loops removed as the search model (PDB ID: 1swt). All models were refined using Refmac520 and model building was carried out with the program Coot. The chemical description of 7-HCAA was taken from Henderson et al.[4] All structural figures were made with the PyMOL molecular graphics software.[5] Structures and all supporting data have been deposited in the Protein Databank.

### Supplemental Schemes and Figures

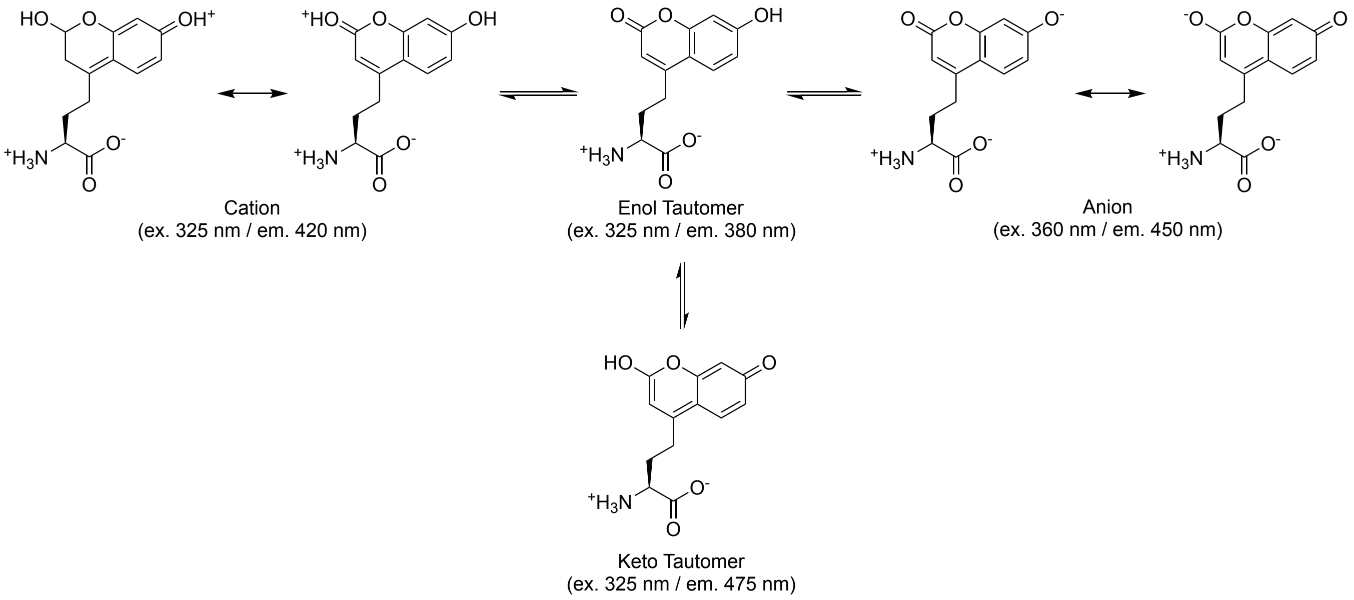


**Scheme S1.** Tautomeric and ionized forms of L-(7-hydroxycoumarin-4-yl)ethylglycine and their excitation/emission wavelengths. Values are from small molecule studies performed in aqueous solutions.[6]

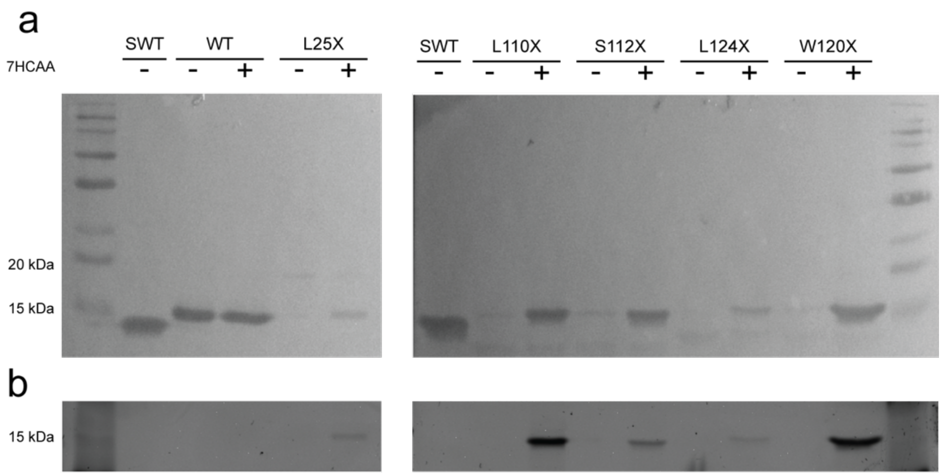


**Figure S2.** Streptavidin mutant expression. (a) Western blot and (b) fluorescent images of whole cell *Escherichia coli* lysates for each streptavidin mutant. Each mutant was expressed in both the presence (+) or absence (-) of 7-HCAA. 1 μg of commercially purchased wild-type streptavidin (SWT) without a 6x-his tag was used as a standard. Faint bands can be detected in the L25X, L110X, S112X, L124X, and W120X lanes without 7-HCAA, which is likely a consequence of amber codon suppression with tyrosine in lieu of 7-HCAA. Faint bands below each of the full-length L110X, S112X, L124X, and W120X bands are likely a consequence of truncated protein in which the amber stop codon was not fully suppressed.

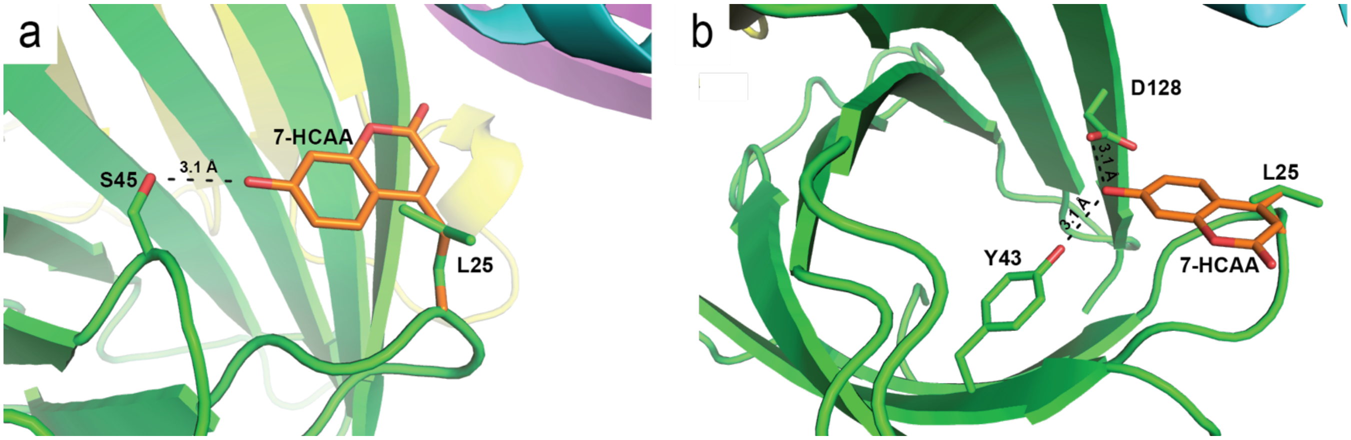


**Figure S3.** Potential interactions between 7-HCAA and surrounding residues in the L25X mutant. 7-HCAA was overlaid on residue L25 using the PyMOL molecular viewing software by fitting the Cα, Cβ, and Cγ atoms of 7-HCAA to the identical atoms in the native leucine. Rotation about $\chi_2$ and $\chi_3$ was carried out in order to simulate conformational sampling that might occur in the mutant protein. A number of potential interactions with surrounding residues were observed, name (a) residue S45 and (b) residues Y43 and D128 in the binding pocket of apo SAV.

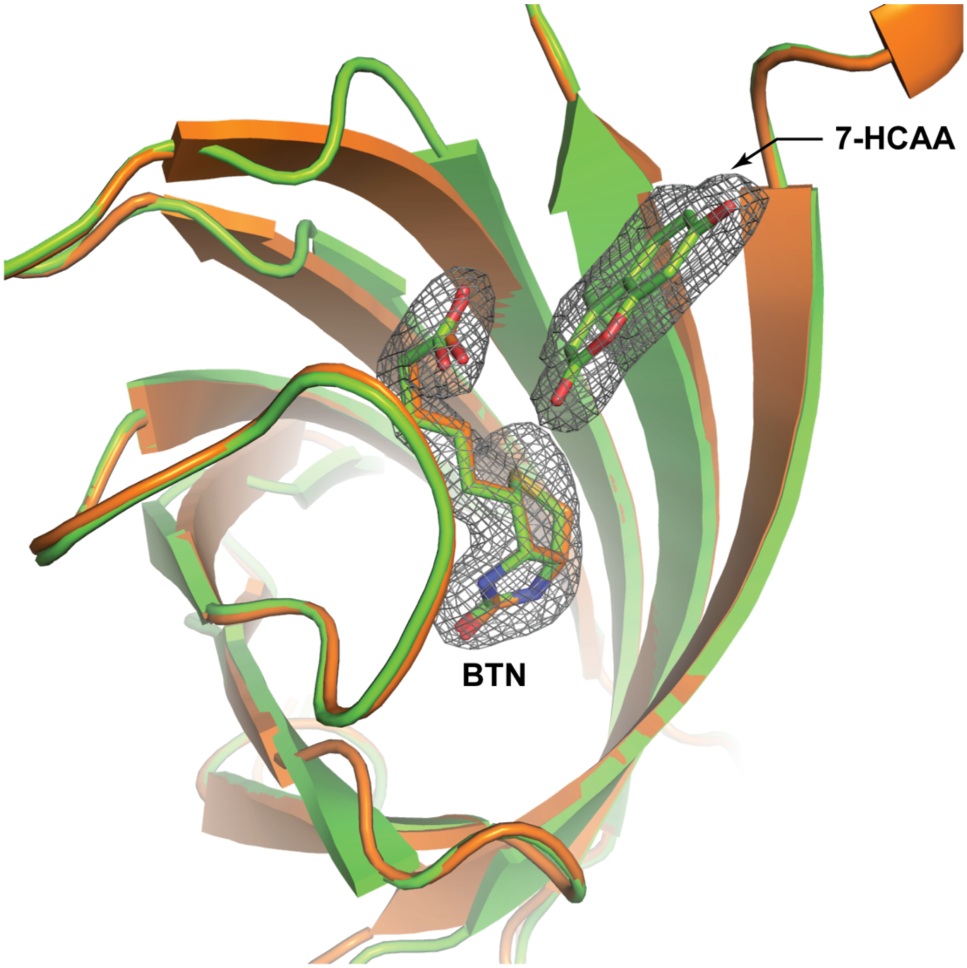


**Figure S4.** Comparison of biotin (BTN) binding in L110X and wt SAV. Holo L110X mutant (green) superimposed with chain A of wild type, biotin-bound SAV (orange, PDB ID: 3ry2). All electron density displayed in this figure derives from the L110X crystal structure. Biotin molecules from the L110X mutant and wild type SAV are shown in green and orange sticks, respectively. Structures were aligned using the super command in PyMOL. The RMSD of the biotin in the L110X crystal structure to the biotin in 3ry2 was calculated to be 0.149 Å using the rms_cur command in PyMOL. Electron density around the L110X biotin is shown as a $2F_O$-$F_C$ map contoured to 1 σ.

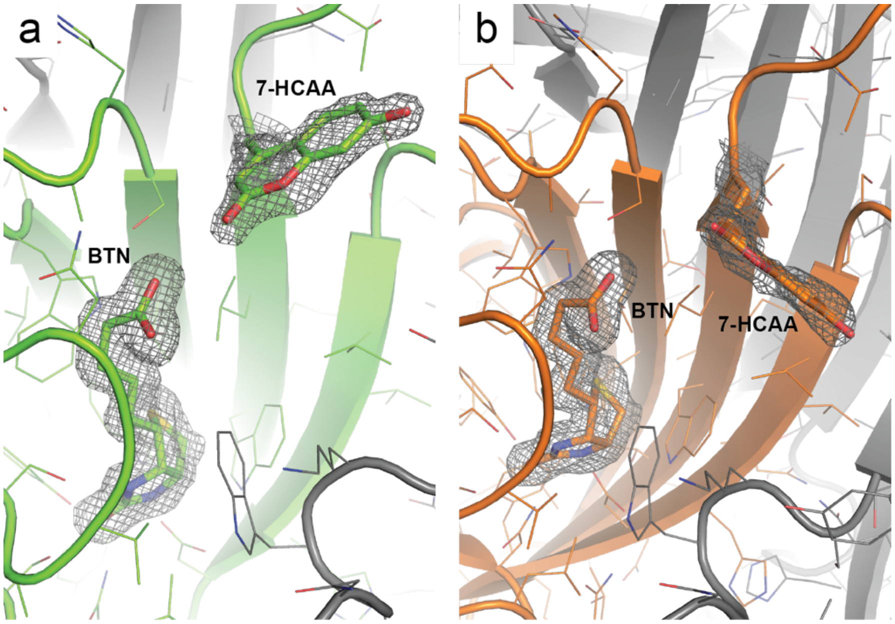


**Figure S5.** Structures of S112X containing biotin (BTN). The orientations adopted by 7-HCAA differ between subunits in this structure. (a) Chain A (green) and (b) chain B (orange) of the asymmetric unit are shown. Symmetry related chains are colored in grey. Electron density around each residue is shown as a $2F_O$-$F_C$ map contoured to 1 σ.

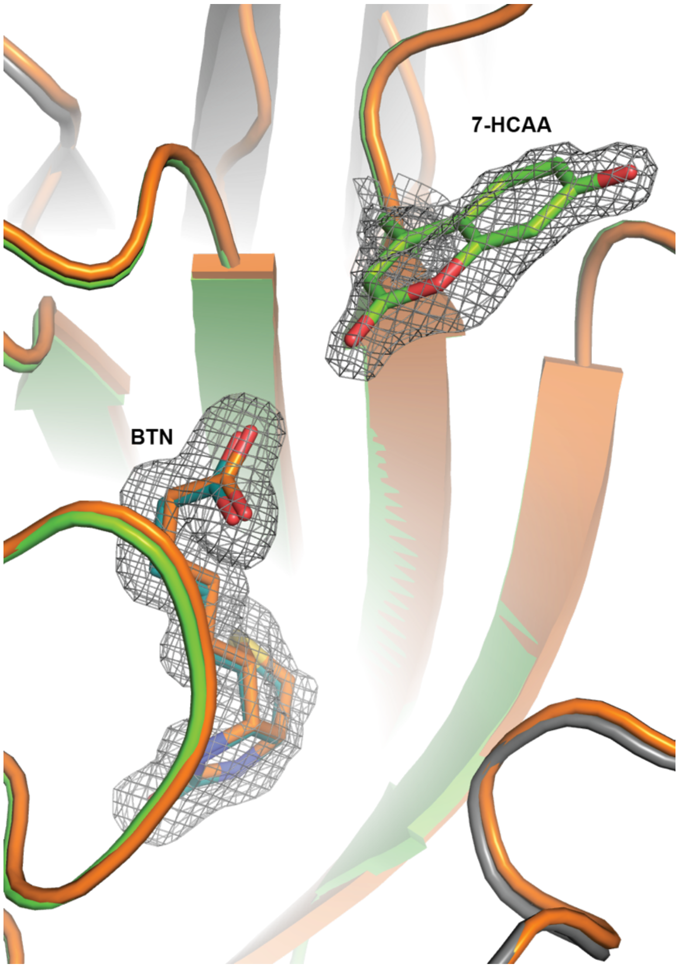


**Figure S6.** The structure of the biotin (BTN) bound S112X mutant (green) is superimposed with biotin bound wild type SAV (orange, PDB ID: 3ry2). Structures were aligned using the super command in PyMOL. The RMSD of the biotin in the S112X crystal structure to the biotin in 3ry2 was calculated to be 0.275 Å using the rms_cur command in PyMOL. Electron density around the S112X biotin is shown as a $2F_O$-$F_C$ map contoured to 1 σ.

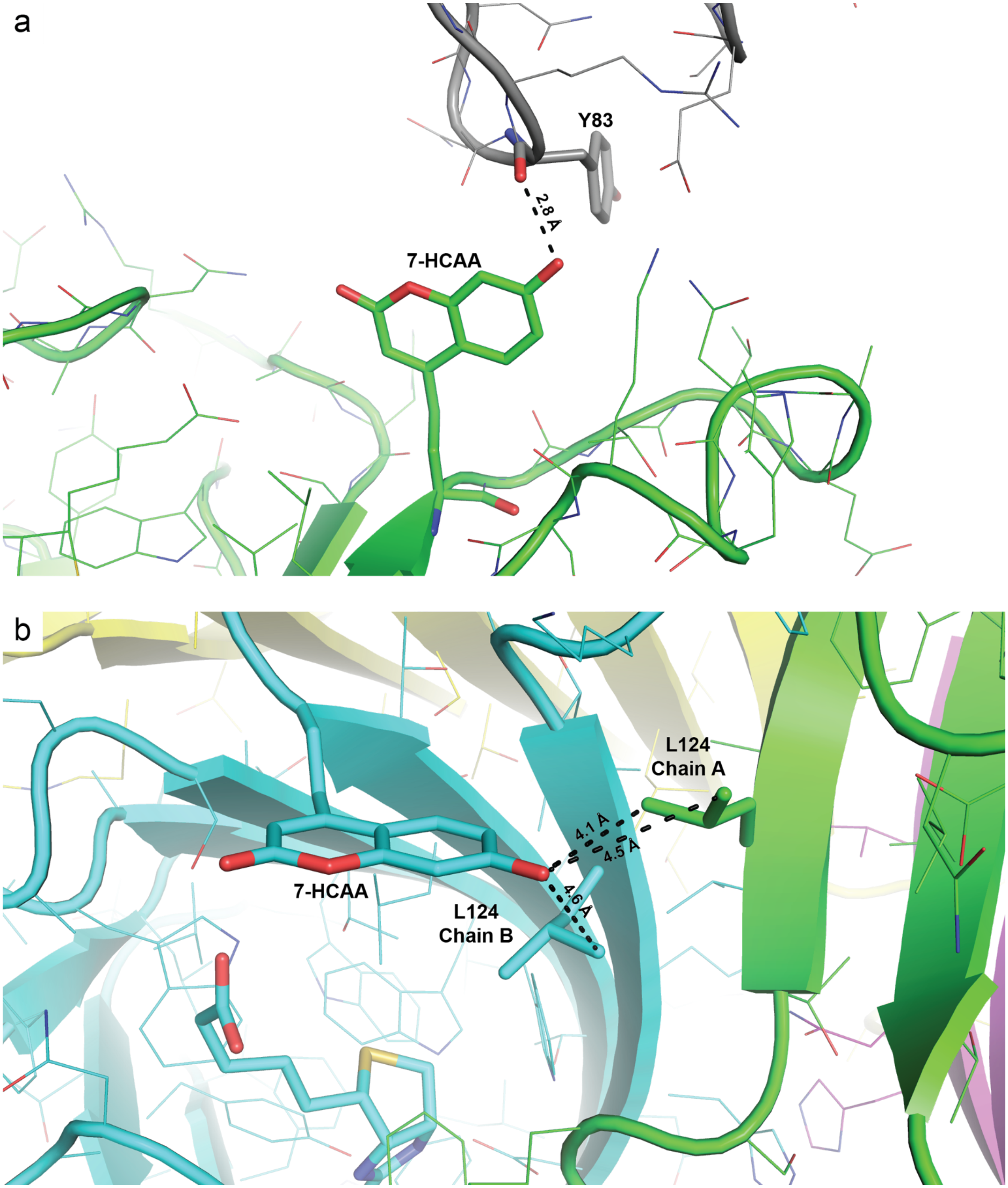


**Figure S7.** Crystal structures of the S112X mutant showing interactions between the 7-HCAA side and surrounding residues. Two conformations of 7-HCAA were observed in chains A (a) and B (b) of this structure. Chain A (a) shows a hydrogen bond between the 7-HCAA and the Y83 of a subunit in an adjacent crystal. In chain B (b), 7-HCAA's phenol is found in in a more hydrophobic environment created by symmetry-related L124 residues in chains A and B (panel B). Dashed black lines illustrate distances between the 7-HCAA phenol and atoms on near neighbor residues; they are not indicative of hydrogen bonds.

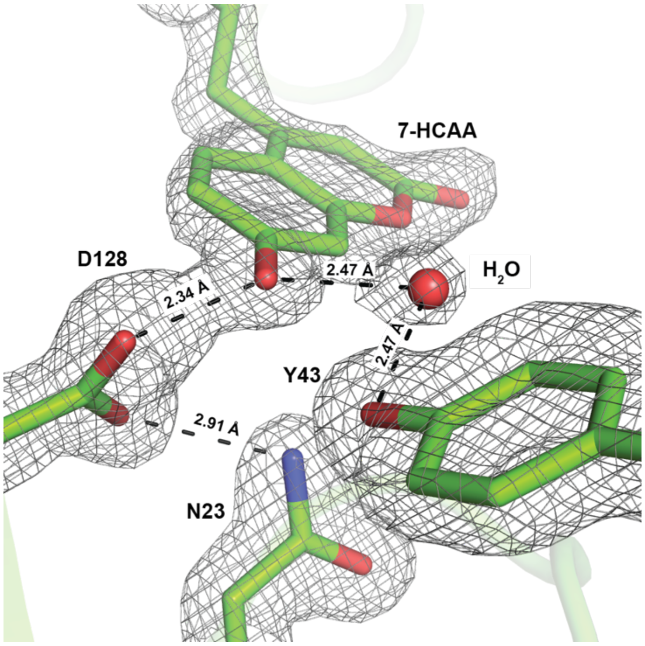


**Figure S8.** An apparent hydrogen bonding network in the W120X mutant. Distances are the averages of the two monomeric streptavidin chains in the asymmetric unit. Electron density around each residue is shown as a $2F_O$-$F_C$ map contoured to 1 σ.

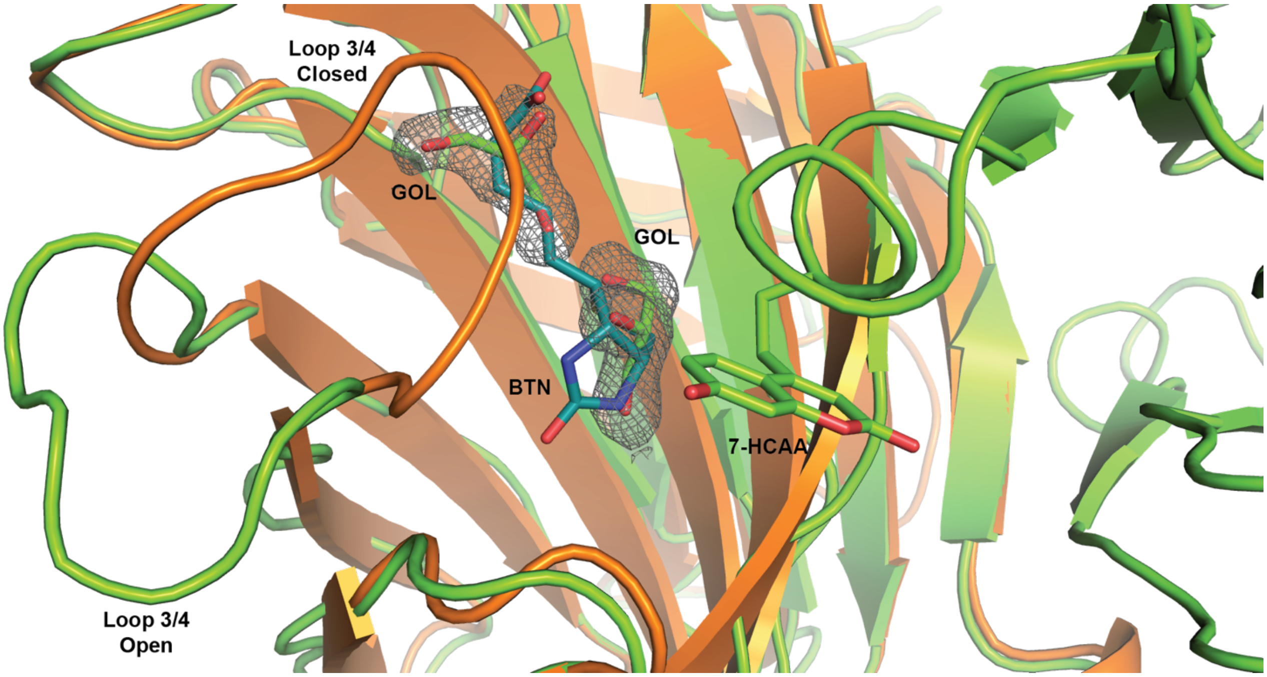


**Figure S9.** A crystal structure of W120X solved when glycerol was used as a cryoprotectant (green) superimposed on a biotin (BTN; teal sticks) bound SAV structure (PDB 3ry2, orange). The bound glycerol (GOL; green sticks) molecules in our structure occupy the biotin binding pocket. Loop 3/4 is also observed in the open position in the W120X crystal structure. The open position of this loop may lead to the observed differences between the W120X + biotin and W120X + glycerol fluorescence spectra shown in figure S10. It is also possible that the biotin itself contributes to these differences. Electron density around each glycerol molecule is shown as a 2FO-FC map contoured to 1 σ.

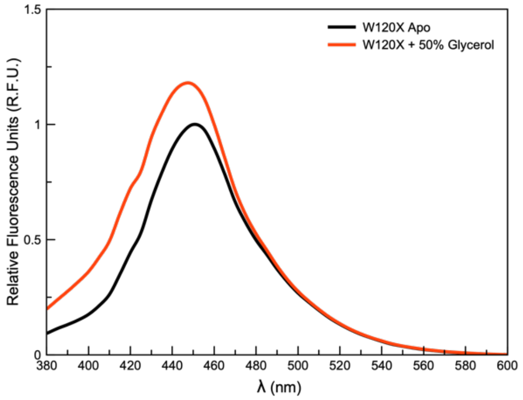


**Figure S10.** A normalized fluorescence spectrum of W120X in the absence (black) and in the presence (red) of glycerol. The protein concentration was ~10 uM and glycerol was present at 50%. An excitation wavelength of 340 nm was used.

**Table S1.** Parameters and statistics for L110X, S112X, and W120X apparent $K_d$ curve fits

| Construct | L110X | S112X | W120X |
|---|---|---|---|
| **Best-fit values** | | | |
| Bottom | 1.83 RFU | 0.4543 RFU | -2.149 RFU |
| Top | 96.62 RFU | 101 RFU | 92.86 RFU |
| $K_d$ | 560.3 nM | 1.61 μM | 6.28 nM |
| HillSlope | -6.857 | 1.12 | 1.301 |
| log $K_d$ | -6.252 | -5.794 | -8.202 |
| Span | 94.79 RFU | 100.5 RFU | 95.01 RFU |
| **95% CI (profile likelihood)** | | | |
| Bottom | -0.8138 to 4.371 RFU | -1.734 to 2.576 RFU | -12.81 to 7.383 RFU |
| Top | 95.11 to 98.96 RFU | 97.36 to 104.9 RFU | 85.12 to 101.4 RFU |
| $K_d$ | 543.0 to 576.8 nM | 1.38 to 1.90 μM | 4.03 to 1.06 nM |
| HillSlope | N/D$^{\alpha}$ | 0.9523 to 1.358 | 0.6759 to 2.567 |
| log $K_d$ | -6.265 to -6.239 | -5.861 to -5.722 | -8.394 to -7.976 |
| **Goodness of Fit** | | | |
| Degrees of Freedom | 26 | 14 | 5 |
| R squared | 0.9947 | 0.9969 | 0.9891 |
| Sum of Squares | 303 | 95.97 | 161.3 |
| Sy.x | 3.414 | 2.618 | 5.68 |
| **Number of points** | | | |
| # of X values | 30 | 18 | 9 |
| # Y values analyzed | 30 | 18 | 9 |

$^{\alpha}$The value was not definable.

**Table S2.** Crystallographic statistics.

| Structure | L110X Apo | L110X Holo | S112X Holo | W120X Apo | W120X Glycerol |
|---|---|---|---|---|---|
| **Data collection** | | | | | |
| Space group | P 1 2 1 1 | P 1 2 1 | I 2 2 2 | I 4 | I 4 1 |
| Cell dimensions<br>a, b, c (Å)<br>α, β, γ (deg) | <br>46.46, 85.56, 58.10<br>90.000, 99.0, 90.00 | <br>50.81, 98.28, 52.65<br>90.00, 112.3, 90.00 | <br>46.209, 93.110, 104.369<br>90.000, 90.000, 90.000 | <br>57.351, 57.351, 172.320<br>90.000, 90.000, 90.000 | <br>57.519, 57.519, 173.026<br>90.000 90.000 90.000 |
| Total Reflections | 614,572 | 340,473 | 36,873 | 956,126 | 2,652,590 |
| Unique Reflections | 64,975 | 27,711 | 19,561 | 40,032 | 44,646 |
| Resolution (Å) | 50.00-1.55<br>(1.58-1.55) | 50.00-2.10<br>(2.14-2.10) | 69.48-1.84<br>(9.03-1.84) | 50.00-1.55<br>(1.58-1.55) | 50.00-1.50<br>(1.53-1.50) |
| I/σ(I) | 10.9 (1.3) | 13.75 (1.9) | 16.7 (5.9) | 32.83 (2.36) | 36.00 (2.14) |
| $R_{meas}$ | 0.062 (0.743) | 0.160 (0.894) | 0.037 (0.187) | 0.115 (1.047) | 0.077 (1.105) |
| $R_{pim}$ | 0.030 (0.364) | 0.076 (0.442) | 0.026 (0.132) | 0.036 (0.341) | 0.021 (0.310) |
| $CC_{1/2}$ | (0.938) | .725 (0.582) | 0.998 (0.951) | 0.999 (0.794) | 1.000 (0.845) |
| $R_{merge}$ | -- | -- | 0.026 (.132) | -- | -- |
| Completeness (%) | 99.6 (93.6) | 97.1 (92.6) | 98.7 (99.2) | 97.9 (99.9) | 100.0 (100.0) |
| Redundancy | 2.1 (2.0) | 2.2 (2.0) | 1.9 (1.9) | 5.2 (4.6) | 6.8 (6.4) |
| **Refinement** | | | | | |
| Resolution (Å) | 42.78-1.55 | 49.14-2.10 | 69.48-1.84 | 43.08-1.55 | 29.65-1.50 |
| $R_{work}$ | 0.166 | 0.194 | 0.174 | 0.173 | 0.126 |
| $R_{free}$ | 0.201 | 0.256 | 0.212 | 0.197 | 0.167 |
| rmsd bond lengths (Å) | 0.0221 | 0.0161 | 0.0197 | 0.0245 | 0.0307 |
| rmsd bond angles (deg) | 2.147 | 1.919 | 2.051 | 2.546 | 2.437 |

*The values in parentheses indicate statistics for the highest resolution shell.
